# Engineering Linear Cap-independent mRNA Vaccines with Intrinsic Adjuvanticity for Potent Cancer Immunotherapy

**DOI:** 10.1101/2025.10.27.684966

**Authors:** Hongwu Yu, Yu Yang, Peng Lin, Chengye Liu, Yifan Wen, Zhuting Fang, Zhixiang Hu, Shenglin Huang

**Affiliations:** Department of Integrative Oncology, Fudan University Shanghai Cancer Center, and Shanghai Key Laboratory of Medical Epigenetics, Institutes of Biomedical Sciences, Fudan University, Shanghai, 200032, China; Department of Oncology, Shanghai Medical College, Fudan University, Shanghai, 200032, China; Department of Oncology and Vascular Interventional Therapy, Clinical Oncology School of Fujian Medical University, Fujian Cancer Hospital (Fujian Branch of Fudan University Shanghai Cancer Center), Fuzhou, 350014, China; State Key Laboratory of Genetics and Development of Complex Phenotypes, Fudan University, Shanghai, China

**Keywords:** mRNA, Cancer immunotherapy, Cancer vaccines, Cap-independent translation, Subgenomic flaviviral RNA

## Abstract

mRNA cancer vaccines have shown promising efficacy in early-phase clinical trials, but existing platforms struggle to boost antitumor efficacy without added cost or complexity. Here, we present a streamlined linear cap-independent RNA (LciRNA) cancer vaccine platform that achieves stable expression *in vivo* without 5’ capping or ribonucleotide modification, and innately stimulates immune responses to enhance antitumor immunity. By fusing a UPA protective sequence, composed of a viral exoribonuclease-resistant RNA (xrRNA) and a poly(A) binding protein (PABP) motif, with an optimized *Enterovirus* A internal ribosome entry site, LciRNA resists 5’ exonuclease decay and drives superior *in vivo* expression. Mechanistically, UPA sequence not only impedes exonuclease-mediated decay but also recruits RNA-binding proteins to stabilize LciRNA. Moreover, LciRNA robustly activates dendritic cell pattern-recognition receptor pathways, promotes dendritic cell maturation, and upregulates proinflammatory signals. In murine melanoma and HPV-associated tumor models, it elicits strong systemic and intra-tumoral T cell responses and superior tumor control, demonstrating how immune stimulation-translation synergy underpins its efficacy. This work establishes a next-generation cost-effective mRNA cancer vaccine platform with simplified production and enhanced efficacy, highlighting immuno-translation coupling as a paradigm for future mRNA cancer vaccines.

## INTRODUCTION

Messenger RNA (mRNA) has emerged as a transformative vaccine platform owing to its rapid development, robust immunogenicity, and favorable safety profile. The successful application of mRNA vaccines during the COVID-19 pandemic demonstrated their exceptional safety and superior protective efficacy compared with traditional modalities^1^. Accordingly, mRNA technology is being actively explored for oncological applications, where it is anticipated to enable the development of a new generation of transformative cancer therapeutics^2,3^. Recent clinical results underscore this promise. In a Phase IIb trial, the personalized vaccine mRNA-4157 plus pembrolizumab reduced the risk of recurrence or death by 44% in advanced melanoma patients^4^. Similarly, the personalized vaccine BNT-122 induced markedly prolonged median recurrence-free survival (RFS) in responders in a Phase I adjuvant trial for pancreatic ductal adenocarcinoma, and elicited long-lived effector T cell clones with an estimated lifespan of 7.7 years^5,6^.

Currently, mRNA platforms for cancer vaccines include conventional linear mRNA, circular RNA (circRNA), and self-amplifying mRNA^7^. Conventional linear mRNA has already been advanced to clinical application and represents the most mature mRNA vaccine technology to date. Wesselhoeft et al. described the circRNA expression platform in 2018, which enables efficient and stable protein expression via RNA circularization and internal ribosome entry site(IRES)-mediated translation^8^. We previously engineered an optimized circRNA vaccine platform specifically for cancer vaccine applications, which demonstrated superior antigen expression and immunogenicity compared to conventional linear mRNA^9^. However, circRNA production requires complex circularization and purification procedures, substantially complicating production workflows and increasing costs^10,11^. The self-amplifying mRNA (saRNA) vaccines, encodes additional viral RNA replication proteins to achieve *in vivo* RNA amplification and robust efficacy at low doses^12^. Nevertheless, because these vaccines produce a large amount of double-stranded RNA (dsRNA) and express multiple heterologous viral proteins, they can induce systemic reactogenicity, potentially compromising tolerability and therapeutic efficacy^13,14^. Furthermore, chemically modified poly(A) mRNA^15^ and multi-capped mRNA^16^ strategies reported by Chen et al. substantially enhance stability and translation but introduce additional manufacturing complexity and tolerability concerns. To date, no existing platform achieves high potency without adding design or production complexity.

By contrast, conventional linear mRNA remains the most readily manufactured vaccine platform, with minimal intrinsic adverse effects ^17^. However, linear mRNA’s optimal translation efficiency and stability requires two essential modifications: 5’ capping^18,19^ and modified ribonucleotides^20,21^. These modifications also increase both production complexity and cost. Circumventing the reliance of linear mRNA on 5’ capping and ribonucleotide modifications could enable the development of a streamlined vaccine platform that balances affordability, manufacturability, and therapeutic efficacy. Studies have explored linear mRNA designs lacking a 5’ cap structure^22–24^, but they generally fail to provide sufficient protection to the mRNA’s 5’ end and, therefore, are unlikely to serve as viable substitutes for traditional capped mRNA.

The mRNA 5’ cap not only recruits eIF4E to facilitate translation initiation but also shields the transcript’s 5’ terminus from XRN-1-mediated exonucleolytic decay ^25,26^, whereas incorporation of modified ribonucleotides enhances mRNA stability^20,21^. A survey of the literature on viral and eukaryotic mRNA metabolism reveals that these functions can, in principle, be conferred entirely by RNA sequences and structural elements. For example, *Flavivirus* genomes harbor structured elements that block XRN-1 to generate abundant subgenomic flaviviral RNAs (sfRNAs) during infection^27,28^, which could be repurposed to shield synthetic transcripts from exonucleolytic digestion. Likewise, certain cellular RNA-binding proteins (RBPs), such as poly(A) binding proteins (PABPs) and IGF2BP1, stabilize mRNAs. Specifically, PABPs binds and stabilizes the poly(A) tail of most mRNAs^29^, whereas IGF2BP1 associates with MYC and several other mRNAs to enhance their half-lifes^30^. Integrating RBP-binding motifs is a viable strategy to further bolster mRNA stability. Finally, IRES, such as our previously characterized EV-A IRES, can efficiently drive cap-independent translation, effectively replacing the 5’ cap’s translational role^8,9^.

In this study, we introduce a novel, entirely sequence-encoded linear cap-independent RNA (LciRNA) platform that requires no 5’ cap or ribonucleotide modifications by integrating the above-mentioned elements. We began by using RNA secondary-structure modeling and luciferase screening to identify an optimal XRN-1-resistant element from flaviviral sfRNA. This element was fused to a PABP-binding sequence and an optimized *Enterovirus* A (EV-A) IRES variant, generating LciRNA constructs that drive robust *in vivo* expression. Mechanistic analyses confirmed that these elements both block XRN-1 decay and recruit endogenous RBPs, including PABP, IGF2BP1, and YBX1, to stabilize transcripts. Additionally, LciRNA vaccines exhibit strong intrinsic adjuvant activity, promoting dendritic cell maturation and proinflammatory cytokine release. Finally, we assessed the safety profile of LciRNA in murine models and formulated LciRNA-based cancer vaccines, which exhibited superior therapeutic efficacy in both B16F10 melanoma and HPV associated tumor models.

## RESULTS

### Engineering LciRNA with Flaviviral xrRNA and PABP Motif for Enhanced Stability and Translation

Previously, we characterized an optimized EV-A IRES (hereafter EV-A-S1) that drives efficient cap-independent translation of target proteins^9^. Because a canonical 5’ cap both recruits eIF4E and protects the 5’ end from XRN-1 mediated degradation^26^, we sought RNA elements that could similarly impede XRN-1 decay for cap-independent LciRNA. *Flavivirus* subgenomic RNAs (sfRNAs) contain structured exoribonuclease-resistant RNA (xrRNA) and dumbbell elements to stall XRN-1 progression^27,31,32^. Moreover, the mRNA 3’ poly(A) tail recruits PABPs, which shield transcripts from the 3’ terminus^29^, suggesting that integrating a PABP-binding motif may also enhance 5’-end protection. We therefore designed LciRNA constructs that incorporate, at the 5’ end, an xrRNA/dumbbell element and a PABP-binding motif upstream of the EV-A-S1 IRES, and, at the 3’ end, a 3’ untranslated region (UTR) followed by a poly(A) tail (Figure 1A). To identify the most effective XRN-1-resistant structure, we retrieved sfRNA sequences of five flaviviruses—Cellular fusion agent virus (CFAV), Dengue virus (DENV), Usutu virus (USUV), Yellow fever virus (YFV), and Zika virus (ZIKV)—from the NCBI virus database, referred to both published sfRNA structural annotations^27^ and RNAfold^33^ predicted secondary structures (Figure S1), yielding 18 candidate xrRNA/dumbbell sequences (Figure 1B).

**Figure 1.**
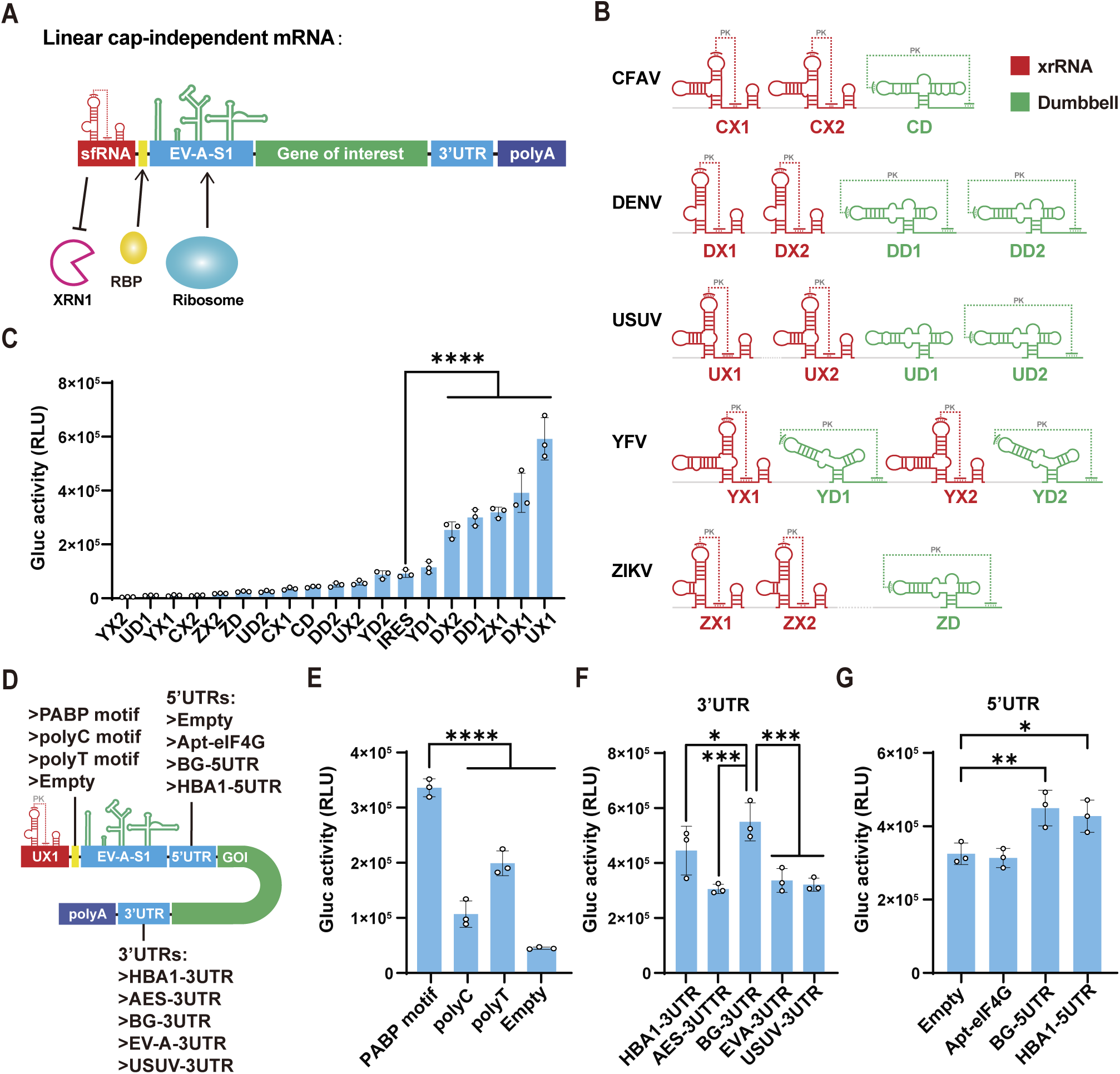
Engineering LciRNA with Flaviviral xrRNA and PABP Motif for Enhanced Stability and Translation. (A) Schematic diagram of LciRNA design. XRN-1 resissting structures of sfRNA and PABP binding motif are placed at the 5’ end to protect LciRNA from degradation, with EV-A-S1 downstream to initiate translation. (B) Schematic diagram of XRN-1 resisting structures in sfRNA of CFAV, DENV, USUV, YFV and ZIKV. xrRNA structures are in red; Dumbbell structures are in green. (C) Comparison of Gluc activity between Gluc LciRNAs with individual structure in (B) or without any XRN-1 resisting structures (Indicated as IRES). (D) Schematic diagram of untranslated regions screened. (E) Comparison of Gluc activity between Gluc LciRNAs with PABP motif, poly cytosine (polyC), poly thymine (polyT) or with no nucleotides (Empty) in the region between UX1 and EV-A-S1. (F) Comparison of Gluc activity between Gluc LciRNAs with indicated 3’UTR. (G) Comparison of Gluc activity between Gluc LciRNAs with indicated 5’UTR or without any nucleotides (Empty). All data are mean (SD) for n= 3 biological replicates. One-way ANOVA was used to calculate the statistical significance. *P < 0.05 was considered statistically significant. **P < 0.01, ***P < 0.001 and ****P < 0.0001 were considered highly significant. ns, not significant.

We inserted each candidate motif at the 5’ end of a Gaussia luciferase (Gluc) reporter LciRNA carrying EV-A-S1 to enable rapid transcript stability screening via luminescence. All constructs were *in vitro* transcribed without cap analogs or modified nucleotides and transfected into HEK293T cells. After 48 h, luciferase assays revealed that UX1, DX1, ZX1, DD1, and DX2 significantly outperformed the unprotected control (Figure 1C). Notably, UX1, derived from USUV, yielded the highest expression and was therefore selected for further experiments.

Because untranslated regions modulate translation in circRNAs and capped mRNAs, we next optimized all other untranslated regions including conventional 5’ and 3’ UTRs in LciRNA (Figure 1D). Incorporating a PABP motif between UX1 and EV-A-S1 yielded higher Gluc expression than polyC motif, polyU motif, or no insert (Empty) (Figure 1E). Screening various 5’ and 3’ UTRs revealed that human β-globin sequences at both sites provided maximal translation (Figure. 1F, 1G). Accordingly, all subsequent LciRNA designs incorporate β-globin 5’ and 3’ UTRs.

### Ribonucleotide Modification and UX1’s Structural Alteration Impair LciRNA Performance

Ribonucleotide modifications often enhance mRNA stability and expression, so we evaluated their impact on LciRNA. Prior studies have shown that N1-methyl-pseudouridine (m1Ψ) improves mRNA stability and translational efficiency^21,34^, m6A enhances circRNA translation^35^, and N4-acetylcytidine (N4-Ac-C) boosts IRES-mediated translation^36^. We thus incorporated 5–100% m6A, m1Ψ, or N4-Ac-C into Gluc LciRNAs. Transfected into HEK293T cells, these modified RNAs exhibited dose-dependent reductions in luciferase activity after 48 h (Figure 2A). This decline likely reflects disruption of critical UX1 and IRES structures by noncanonical bases^37^, underscoring that LciRNA’s function relies on precise folding and is incompatible with extensive ribonucleotide modifications.

**Figure 2.**
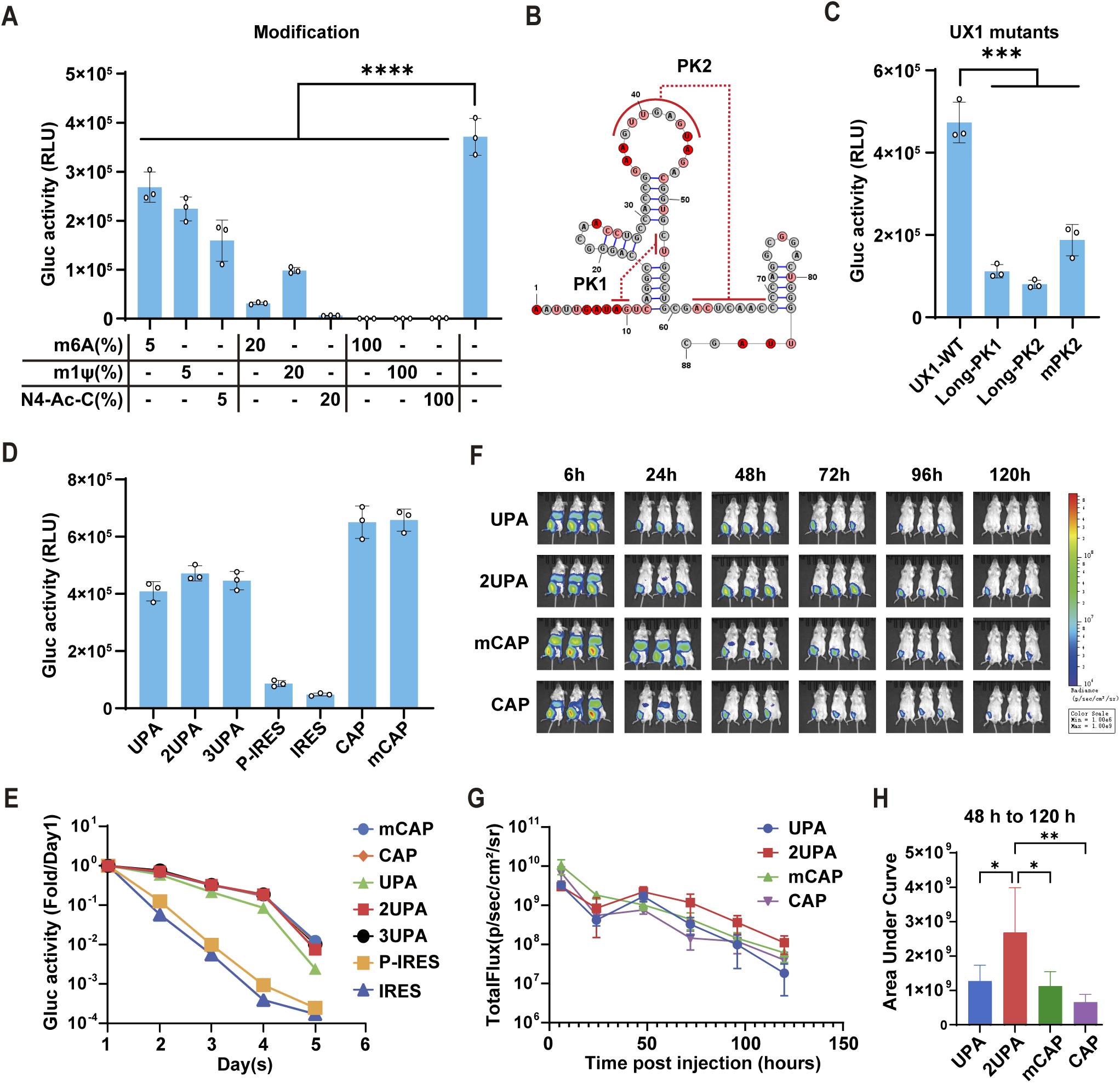
Engineered LciRNA Outperforms m1Ψ-modified Capped mRNA in *in vivo* Persistence. (A) Comparison of Gluc activity between Gluc LciRNAs with indicated percentages of m6A, m1Ψ and N4-Ac-C modified nucleotides. (B) Secondary structure of UX1. The nucleotide coloring indicates normalized reactivity values of <=0.4(gray), >0.4 and <0.85 (light red), and >=0.85 (dark red). Bold black characters indicate pseudoknots. (C) Comparison of Gluc activity between Gluc LciRNAs with indicated UX1 mutants or wildtype UX1. (D) Comparison of Gluc activity between Gluc LciRNAs with indicated 5’ end protective sequences (UPA, 2UPA, 3UPA, P-IRES (with only PABP motif) and IRES), capped Gluc mRNA (CAP) and 100% m1Ψ modified capped mRNA (mCAP). (E) Relative Gluc activity of indicated RNAs from day 1 to 5. (F) *In vivo* fluorescence images of mice at indicated time after injection of indicated Fluc RNA-LNP. (G) Total Flux data corresponding to the images in (F). (H) Area under curve analysis of Total Flux data from 48 h to 120 h in (G). All data are mean (SD) for n= 3 biological replicates. One-way ANOVA was used to calculate the statistical significance. *P < 0.05 was considered statistically significant. **P < 0.01, ***P < 0.001 and ****P < 0.0001 were considered highly significant. ns, not significant.

We subsequently examined whether UX1 structural modification could further enhance LciRNA protection. Previous studies demonstrated that xrRNA blocking of XRN-1 is regulated by pseudoknot length^38^. We therefore modeled UX1’s structure using SHAPE-MaP (selective 2’-hydroxyl acylation analyzed by primer extension and mutational profiling) and IPknot analysis, revealing two pseudoknots, PK1 and PK2 (Figure 2B, S2A). Then, we generated UX1 mutants with longer pseudoknots (Long-PK1, Long-PK2) or increased GC content in PK2 (mPK2) and assessed their performance in Gluc LciRNA. Unexpectedly, all three mutants showed significantly diminished expression levels compared with wild-type UX1 (Figure 2C), indicating that the native 88-nt UX1 sequence already achieves optimal XRN-1 resistance.

### Engineered LciRNA Outperforms m1Ψ-modified Capped mRNA in *in vivo* Persistence

Building on our initial observation that coupling UX1 with a PABP-binding motif maximizes LciRNA expression, we designated their combination the UPA protective sequence. We hypothesized that tandem repeats of UPA could further enhance transcript stability. Accordingly, we designed Gluc-encoding LciRNAs featuring one (UPA), two (2UPA), or three (3UPA) tandem UPA units at the 5’ terminus. These constructs were evaluated alongside unmodified (CAP) and m1Ψ-modified (mCAP) Cap1-capped Gluc mRNAs, as well as uncapped RNAs containing only a 5’ PABP motif sequence (P-IRES) or the IRES alone. UPA-containing transcripts exhibited significantly increased luciferase activity, although their expression at 48 h post-transfection remained slightly below that of CleanCap-capped Cap1 mRNAs (Figure 2D). Notably, the 2UPA-LciRNA achieved the highest expression, 10.5-fold higher than IRES-only control, demonstrating that dual UPA repeats confer optimal protection.

To evaluate transcript expression dynamics, we transfected HeLa cells and sampled supernatants daily for five days. The 2UPA and 3UPA LciRNAs sustained stable expression at levels equivalent to capped mRNAs (Figure 2E). In contrast, single UPA LciRNA showed a modest decline, and the IRES-only control dropped by over 10-fold each day; the PABP-IRES LciRNA showed only slight improvement over the unprotected control (Figure 2E). These data confirm that the UPA motif confers robust protection and that tandem UPA repeats restore cellular stability to match that of Cap1-capped transcripts.

To evaluate *in vivo* expression, BALB/c mice were injected intramuscularly with SM102 LNP-encapsulated UPA and 2UPA LciRNA encoding firefly luciferase (Fluc) as well as CAP and mCAP Fluc mRNAs. Bioluminescence imaging over 120 h post-injection revealed that although 2UPA LciRNA Fluc yielded lower luminescence within the first 24 h, its signal surpassed both mCAP Fluc and CAP Fluc from 48 h to 120 h (Figure 2F, 2G, 2H). These findings demonstrate that 2UPA LciRNA confers greater *in vivo* transcript stability than m1Ψ-modified capped mRNA. This *in vivo* profile contrasts with our cell-culture results and underscores differences in mRNA translation and turnover between immortalized cell lines and mammalian tissues. These differences are likely driven by continuous proliferation and homogeneity in immortalized cells. Given that LciRNA vaccines are intended for clinical application, *in vivo* expression kinetics may more accurately predict therapeutic efficacy. In summary, our engineered LciRNA demonstrates superior *in vivo* expression stability compared to m1Ψ-modified Cap1-capped mRNA, suggesting advantages for cancer vaccine applications.

### UPA Sequence Stabilizes LciRNA Through XRN-1 Inhibition and RBP Binding

The above results showed that incorporation of the UPA sequence endows uncapped LciRNA with stability comparable to that of Cap1-capped mRNA. To elucidate the mechanism underlying UPA-mediated stabilization, we assessed its capacity to inhibit XRN-1 mediated degradation and to recruit intracellular RB<u>P</u>s. In an *in vitro* XRN-1 degradation assay (Figure 3A), the uncapped and unprotected RNA (IRES) was completely degraded after 2 h of XRN-1 treatment, whereas UPA and 2UPA LciRNAs as well as capped mRNA (CAP) remained largely intact (Figure 3B). Consistent with Gluc expression results, 2UPA conferred stronger protection than a single UPA, demonstrating that UPA impedes XRN-1 digestion of LciRNA and that tandem duplication further potentiates this effect.

**Figure 3.**
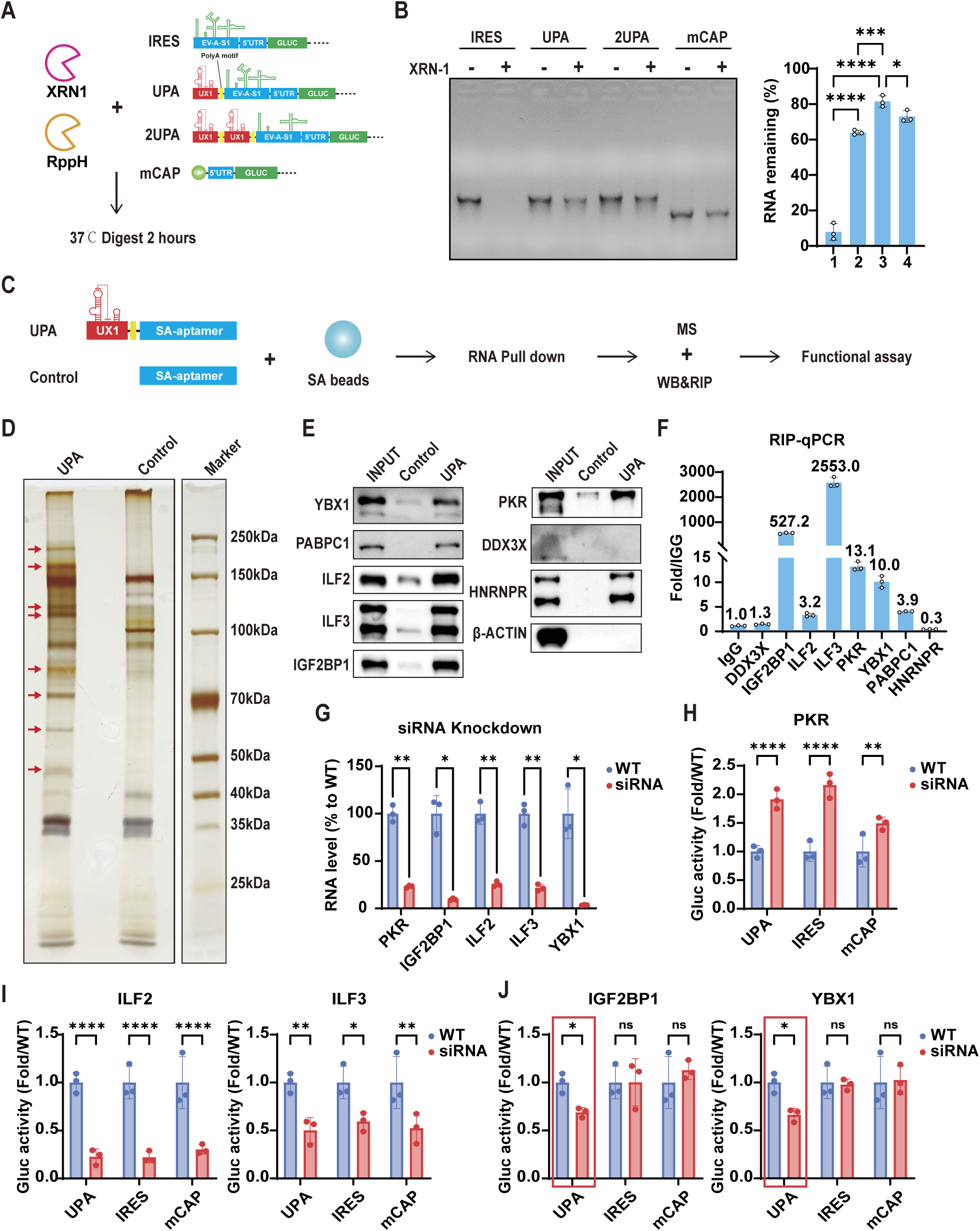
UPA Sequence Stabilizes LciRNA Through XRN-1 Inhibition and RBP Binding. (A) Schematic diagram of XRN-1 *in vitro* digestion assay. XRN-1 and RppH were added together to digest the indicated RNAs. (B) 2% agarose gel analysis of RNAs after 2 hours of XRN-1 digestion. right panel: representative gel image; left panel: quantification of three independent experiments. (C) Schematic diagram of experimental procedures to identify UPA specific RBPs. Briefly, uncapped RNA with UPA plus SA-aptamer sequences or SA-aptamer alone were incubated with streptavidin beads to pull-down interacting proteins, after which mass spectrometry, western blot, RNA immunoprecipitation analysis and functional assays were conducted to verify RBPs’ binding and function. (D) Silver-stained gel of proteins isolated by RNA pull-down assay. Red arrows highlight UPA specific bands. (E) Western blot validation results of potential UPA sequence binding proteins. (F) RNA immunoprecipitation-qPCR (RIP-qPCR) results of potential UPA-binding proteins in Hela cells. Numbers above the bars indicate folds of UPA RNA levels to that of IgG control group. (G) Relative RNA levels quantified by qPCR after siRNA-mediated knockdown of the corresponding genes. Hela cells were treated with siRNAs targeting specific genes for 48 hours prior to qPCR analysis. (H), (I), (J) Relative Gluc activity of three types of GLUC RNAs in Hela cells with siRNA knockdown of the corresponding genes, compared to non-knockdown cells. Hela cells were treated with siRNAs targeting specific genes for 48 hours, followed by transfection with UPA LciRNA, LciRNA containing only IRES, or 100% m1Ψ modified capped mRNA (mCAP). GLUC activity was assessed 24 hours post-transfection. All data are mean (SD) for n = 3 biological replicates. For (B) One-way ANOVA was used to calculate the statistical significance. For (G)-(J) two-tailed unpaired Student’s t-test was used to calculate the statistical significance. *P < 0.05 was considered statistically significant. **P < 0.01, ***P < 0.001 and ****P < 0.0001 were considered highly significant. ns, not significant.

We next sought UPA-binding proteins by fusing the UPA sequence to a streptavidin aptamer (SA)^39^ and performing RNA pull-down with streptavidin-coated beads(Figure 3C). HeLa cell lysates were incubated with beads bearing either the UPA–SA aptamer fusion or SA aptamer alone, and bound proteins were resolved by SDS–PAGE (Figure 3D). Mass spectrometry, western blotting, and RNA immunoprecipitation analyses identified YBX1, PABPC1, ILF2, ILF3, IGF2BP1, and PKR as UPA-specific interactors in the cytoplasm (Figure 3E, 3F, S2B). Notably, ILF2, ILF3, and PKR are immune regulatory proteins, suggesting UPA’s role in immune signaling ^40–44^. PABPC1, which binds poly(A) tails to stabilize transcripts and enhance translation, likely reinforces LciRNA stability and translational efficiency via the PABP motif in UPA. Similarly, YBX1 and IGF2BP1 are well-established RBPs that promote mRNA stability ^30,45,46^, and their association with UPA may further elevate LciRNA stability.

To assess each RBP’s role in LciRNA expression, we performed siRNA-mediated knockdown (excluding PABPC1 to preserve 3’ tail integrity) in HeLa cells (Fugure 3G). We then transfected UPA-Gluc LciRNA, IRES-Gluc RNA, and capped Gluc mRNA and measured Gluc activity. PKR depletion significantly increased expression of all three RNA constructs (Fugure 3H), likely due to relief of PKR mediated eIF2α phosphorylation and global translational repression^41,43,47^. In contrast, knockdown of ILF2 or ILF3 reduced expression of all three RNAs (Fugure 3I), consistent with their reported roles in mitosis and effects on cell viability^48,49^. Notably, silencing IGF2BP1 or YBX1 preferentially decreased UPA-LciRNA expression without affecting the other RNAs (Figure 3J). Given the established RNA-stabilizing functions of IGF2BP1 and YBX1^30,45,46^, these findings indicate that the UPA motif enhances LciRNA stability not only by blocking XRN-1 mediated decay but also via recruitment of IGF2BP1, YBX1, and PABPC1.

### Self-Adjuvanting LciRNA Drives PRR-Mediated DC Maturation with Minimal Toxicity

The ultimate goal of a cancer vaccine is to elicit potent, tumor-specific immune responses, which require both efficient antigen expression and the vaccine’s intrinsic or adjuvant mediated immunostimulatory capacity^50,51^. Unlike conventional capped mRNA, LciRNA possesses a 5’-triphosphate moiety that is recognized by pattern recognition receptors such as RIG-I, thereby potently triggering inflammatory signaling ^52^. Its viral-derived IRES contains extended stem-loop structures that may be sensed by protein kinase R (PKR) and toll-like receptors (TLRs), activating downstream innate immune cascades^53–55^. Furthermore, the UPA’s interactions with PKR, ILF2 and ILF3 potentially enhance cellular recognition of LciRNA^40–44,56^. Collectively, these features suggest that LciRNA engages multiple pattern recognition receptors (PRRs) to initiate innate immune pathways, conferring a self-adjuvant effect that can amplify adaptive antitumor immunity.

To assess the potential immunostimulatory ability of LciRNA vaccines, mouse bone marrow derived dendritic cells (BMDCs) were transfected with Fluc-encoding LciRNAs (UPA-Fluc, 2UPA-Fluc) or capped Fluc mRNAs (CAP-Fluc, mCAP-Fluc). At 24 h post-transfection, total RNA was harvested for quantitative RT-PCR and RNA sequencing. qRT-PCR analysis revealed that BMDCs transfected with either LciRNA construct upregulated MHC-I, the co-stimulatory molecules CD40, CD80, and CD86, and the proinflammatory cytokines IFN-α, IFN-β, and IL-6, to a significantly greater extent than cells receiving capped mRNA; only unmodified CAP-Fluc induced modest increases in CD40, IL-6, and IFN-β (Figure 4A). Transcriptomic analysis demonstrated that LciRNA strongly activated gene sets associated with antigen processing and presentation, RIG-I and TLR signaling, type I interferon response, and inflammatory cytokine production (Figure 4B, 4C). Further examination of PRR-induced dendritic cell (DC) maturation gene signatures^57^ revealed near-complete concordance between genes upregulated by LciRNA and canonical PRR-upregulated genes (Figure S3A), indicating that LciRNA drives DC maturation via PRR engagement. Moreover, LciRNA markedly increased the expression of T cell co-stimulatory molecules, proinflammatory chemokines (Cxcl9, Cxcl10), and cytokines (Il6, Il12, Il15, Ifna1, Ifnb1) (Figure S3B). Collectively, these data confirm that LciRNA potently activates PRR signaling, promotes DC maturation, and induces proinflammatory chemokine and cytokine expression, supporting its strong self-adjuvant capacity to enhance antitumor immune responses.

**Figure 4.**
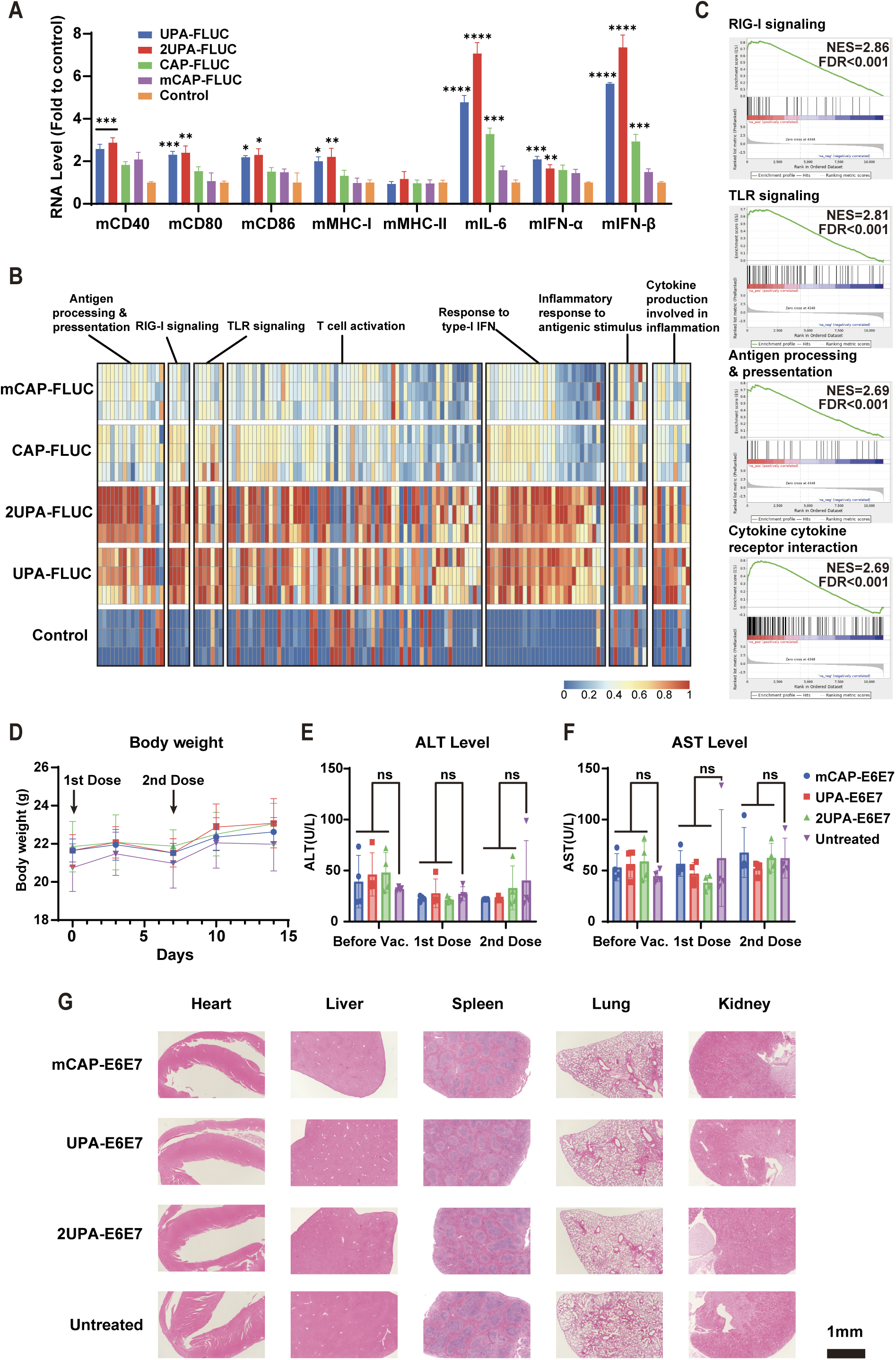
Self-Adjuvanting LciRNA Drives PRR-Mediated DC Maturation with Minimal Toxicity. (A) Normalized RNA levels of BMDCs transfected with indicated LciRNAs (UPA-Fluc, 2UPA-Fluc), capped mRNAs (CAP-Fluc, 100% m1Ψ modified mCAP-Fluc) or empty transfection reagent (Control) for 24 hours. Data were measured by RT-qPCR. The statistical significance was assessed between Control and indicated groups. (B) Heatmap of normalized mRNA expression levels of differentially expressed genes involved in antigen processing & presentation, RIG-1 signaling, TLR signaling, T cell activation, response to type-I interferon, inflammatory response to antigenic stimulus and cytokine production involved in inflammation. Columns represent individual genes and rows represent experimental samples (n = 3 biological replicates per group). TPM values were scaled by column to standardize gene-wise expression patterns, with color intensity indicating relative expression levels (blue, low expression; red, high expression). (C) GSEA plots showing the most-enriched immune response related KEGG pathways in the 2UPA group. FDR stands for false discovery rate. NES stands for normalized enrichment score. (D) Body weight of mice after intramuscular injection of UPA-E6E7, 2UPA-E6E7, or mCAP-E6E7 vaccines. (2 doses 5 μg per dose, black arrow indicated injection time). (E) & (F) Serum ALT (E), AST(F) levels in mice before and after administration of two vaccine doses. (“Before Vac.” indicates before vaccination, while “1st dose” and “2nd dose” means 7 days after the first and second doses, respectively.) (G) Histological staining of major organs two weeks after the second vaccine dose (scale bar = 1mm). All data are mean (SD) for n = 3 biological replicates. One-way ANOVA was used to calculate the statistical significance. *P < 0.05 was considered statistically significant. **P < 0.01, ***P < 0.001 and ****P < 0.0001 were considered highly significant. ns, not significant.

To evaluate potential *in vivo* toxicity of LciRNA vaccines, we formulated SM102 LNPs encapsulating UPA-E6E7 LciRNA, 2UPA-E6E7 LciRNA, or m1Ψ-modified CleanCap-E6E7 mRNA, and administered 5 μg RNA intramuscularly to C57BL/6 mice, followed by a booster dose one week later. Body weight and serum alanine aminotransferase (ALT) and aspartate aminotransferase (AST) levels were measured before immunization and seven days after each dose. No vaccinated group exhibited significant deviations in body weight compared with untreated controls (Figure 4D). Similarly, ALT and AST levels remained comparable to baseline and those of untreated controls (Figure 4E, 4F). Two weeks after the booster dose, histopathological examination of heart, liver, spleen, lung, and kidney via H&E staining revealed no vaccine-related lesions (Figure 4G). These findings demonstrate that LciRNA vaccines are well tolerated in mice at a dosing regimen equivalent to 1 mg per dose in humans.

### LciRNA Vaccination Provokes Potent Tumor-Specific T Cell Responses and Controls B16F10-OVA Melanoma Growth

To evaluate therapeutic efficacy of LciRNA-based cancer vaccines, we established an ovalbumin-expressing B16F10 (B16F10-OVA) melanoma model in C57BL/6 mice. On day 3 and 10 post tumor inoculation, mice received intramuscular doses of 3 μg UPA-OVA LciRNA, 2UPA-OVA LciRNA, m1Ψ-modified capped mCAP-OVA mRNA, or m1Ψ-modified capped Fluc control mRNA (Figure 5A). Tumor growth monitoring revealed that all OVA-targeting vaccines significantly inhibited tumor growth compared with mCAP-Fluc, with the 2UPA-OVA cohort exhibiting the most pronounced inhibition (Figure 5B). On day 20, excised tumors from UPA-OVA and 2UPA-OVA groups were markedly smaller, and complete tumor regression occurred in 3/6 mice per group, versus 1/6 in the mCAP-OVA group (Figure 5C, S4B). Body weights remained stable across all groups, indicating favorable tolerability (Figure S4A). These results demonstrate that LciRNA vaccines provide superior antitumor efficacy over m1Ψ-modified capped mRNA in mouse melanoma model.

**Figure 5.**
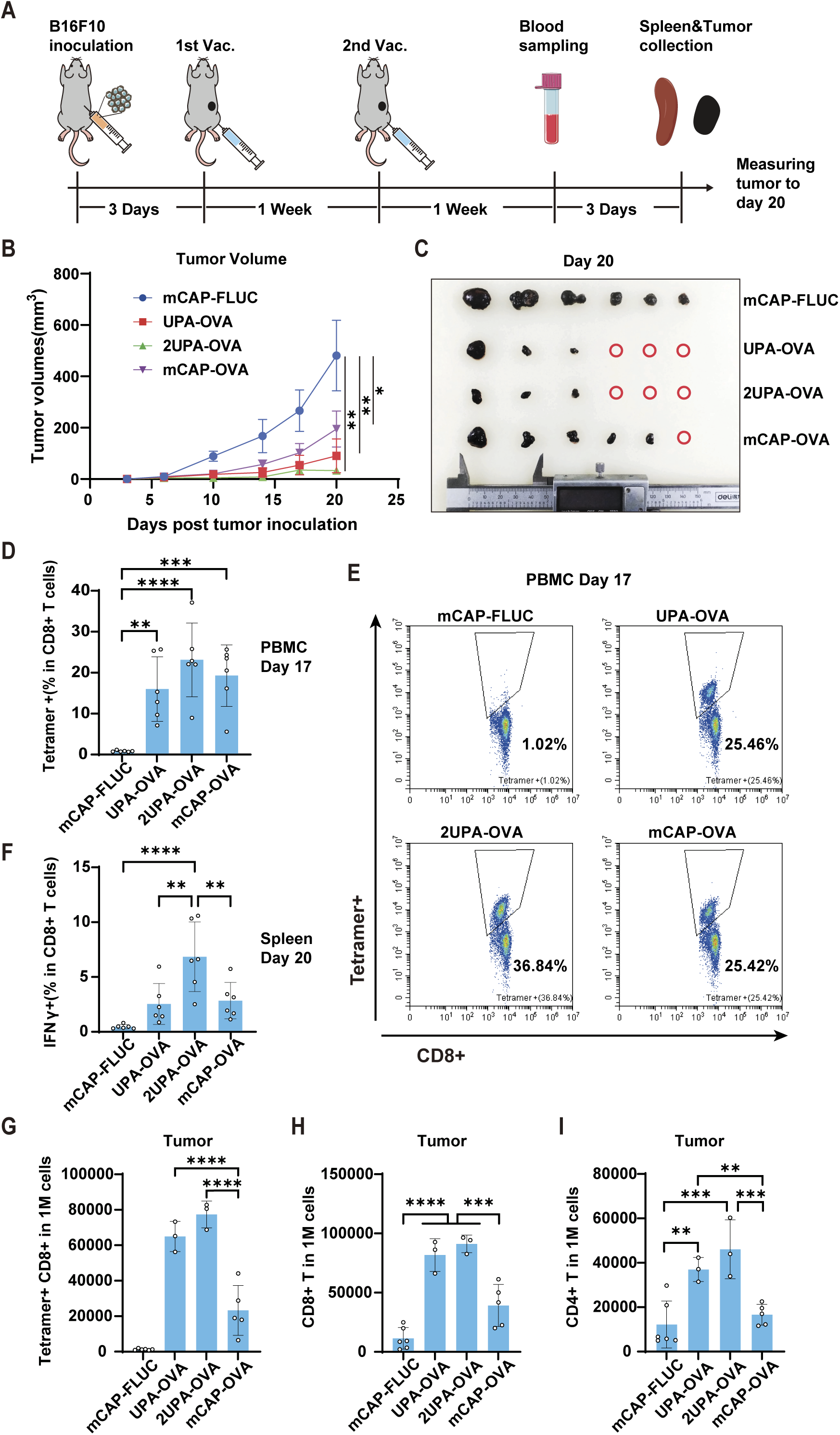
LciRNA Vaccination Provokes Potent Tumor-Specific T Cell Responses and Inhibits B16F10-OVA Melanoma Growth. (A) Timeline of tumor inoculation, vaccination, and blood and tissue sample collection. Mice were subcutaneously inoculated with 1×10^5^ B16F10-OVA cells and intramuscularly injected with mCAP-Fluc, UPA-OVA, 2UPA-OVA, or mCAP-OVA vaccines (3 μg per mouse, n = 6 for each group) on day 3 and day 10. (B) Tumor volumes from day 0 to day 20 after tumor inoculation. Statistical significance was calculated based on tumor volumes data on day 20. (C) Photographs of tumors excised after euthanasia on day 20 post-tumor inoculation, with red circles indicating complete tumor regression in mice. (D) Proportion of OVA-specific CD8+ T cells in PBMCs one week after the second vaccine dose (OVA peptide MHC tetramer-positive). (E) Representative flow cytometry plots for (D). (F) Proportion of OVA-specific CD8+ T cells in the spleen 10 days after the second vaccine dose (IFN-γ-positive T cells after transfection with OVA mRNA). (G) Number of OVA-specific CD8+ T cells per million cells in tumors 10 days after the second vaccine dose (OVA peptide MHC tetramer-positive). (H) Number of CD4+ T cells per million cells in tumor tissue. (I) Number of CD8+ T cells per million cells in tumor tissue. For (B), data are mean (SEM). All other data are mean (SD) for at least 3 biological replicates. One-way ANOVA was used to calculate the statistical significance. *P < 0.05 was considered statistically significant. **P < 0.01, ***P < 0.001 and ****P < 0.0001 were considered highly significant. ns, not significant.

To evaluate antigen-specific T cell responses, peripheral blood mononuclear cells (PBMCs) were collected one week after the booster and stained with OVA (SIINFEKL) Tetramer. FACS analysis revealed that UPA-OVA, 2UPA-OVA, and mCAP-OVA vaccines induced 15.8%, 22.9%, and 19.1% OVA-specific CD8+ T cells on average, respectively, versus 0.7% in control group (Figure 5D, 5E), with 2UPA-OVA eliciting the strongest response. On day 20, spleens and tumors were harvested to further characterize systematic and intratumoral T cell responses. Splenocytes restimulated ex vivo with OVA mRNA exhibited the highest frequency of IFN-γ⁺ CD8⁺ T cells in the 2UPA-OVA group compared to UPA-OVA and mCAP-OVA (Figure 5F, S4C). Analysis of tumor-infiltrating lymphocytes showed that both UPA-OVA and 2UPA-OVA significantly increased the proportion of OVA-specific CD8+ T cells (Figure 5G) and total CD4+ and CD8+ T cell infiltration (Figure 5H, 5I) relative to mCAP-OVA and control. These findings suggest that LciRNA vaccines convert immunologically “cold” tumors into “hot” tumors by enhancing T cell infiltration. Collectively, the 2UPA-OVA LciRNA vaccine demonstrated the most robust antitumor efficacy and T cell induction, while UPA-OVA matched mCAP-OVA in systematic T cell response but achieved greater tumor control, presumably owing to its self-adjuvant activity that enhanced T cell infiltration.

### Effective Clearance of HPV associated Tumors by LciRNA-Based Therapeutic Vaccination

Human papillomavirus (HPV) infection is etiologically linked to cervical and oropharyngeal cancers, in which tumor cells commonly express the viral oncoproteins E6 and E7^58,59^. Therefore, the E6 and E7 proteins serve as ideal target antigens for HPV-associated malignancies. To assess the therapeutic potential of our HPV E6E7 LciRNA vaccines, we established an HPV-associated tumor model by subcutaneously injecting C57BL/6 mice with HPV-E6/E7–expressing TC-1 cells. When tumors reached ∼50 mm³ on day 9, mice received intramuscular injections of 3 μg UPA-E6E7 LciRNA, 2UPA-E6E7 LciRNA, m1Ψ-modified capped mCAP-E6E7 mRNA, or m1Ψ-modified capped Fluc control mRNA (mCAP-Fluc) (Figure 6A). Following two immunizations, all three E6E7 vaccine groups exhibited continuous tumor regression, whereas tumors in control mice progressed rapidly (Figure 6B, S6C). On day 23, mice were photographed, showing significantly smaller tumors in the UPA-E6E7 and 2UPA-E6E7 groups (Figure S6B). Survival analysis demonstrated that all mice in the three E6E7 vaccine groups survived beyond the 40-day observation period, while complete tumor regression was achieved by day 33 in mice vaccinated with UPA-E6E7 or 2UPA-E6E7, compared to day 40 for mCAP-E6E7 (Figure 6C, S6C). These results indicate that the LciRNA vaccines achieved more rapid tumor eradication than m1Ψ-modified capped mRNA.

**Figure 6.**
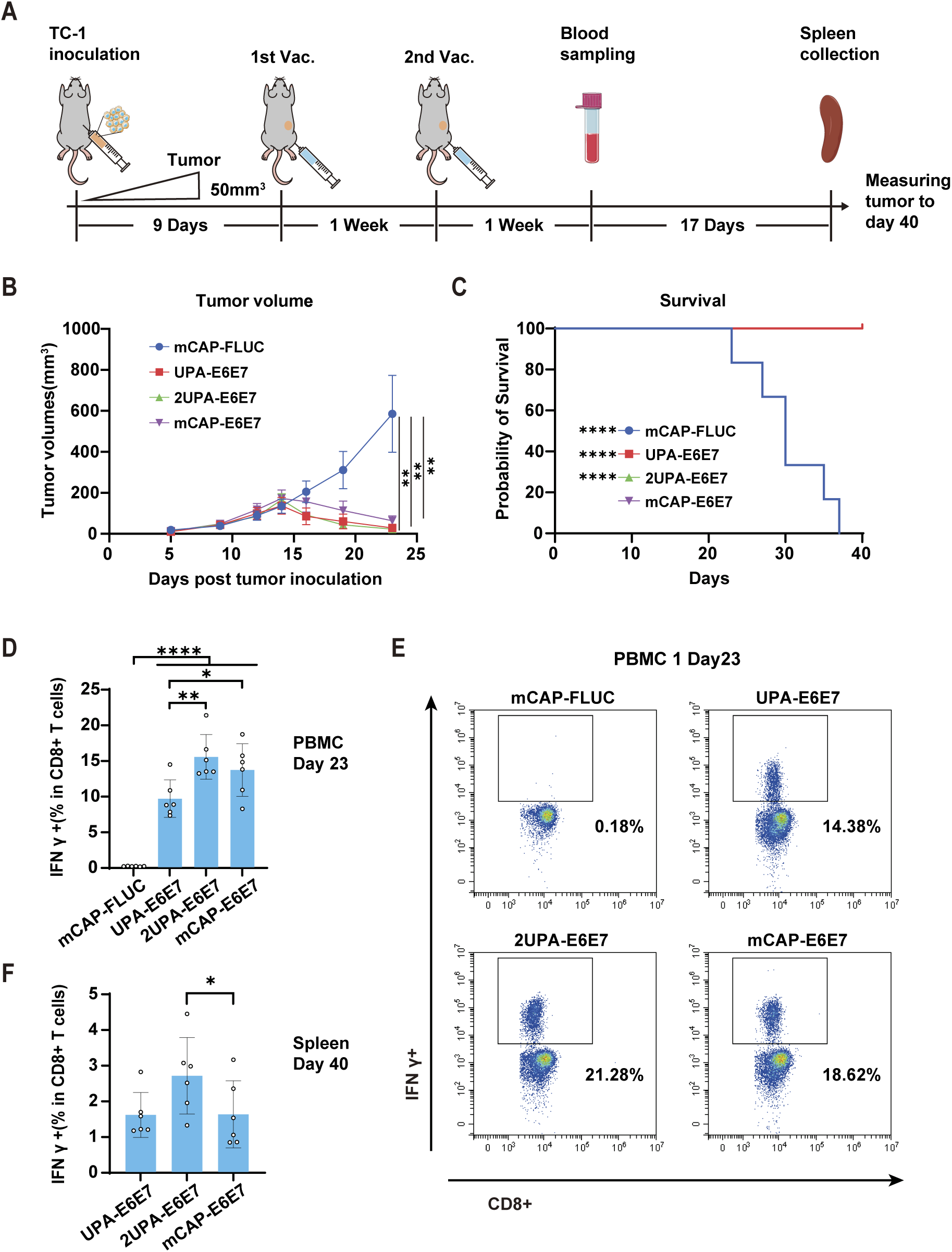
Effective Clearance of HPV associated Tumors by LciRNA-Based Therapeutic Vaccination. (A) Timeline of tumor inoculation, vaccination, and blood and tissue sample collection. Mice were subcutaneously inoculated with 5×10^5^ TC-1 cells, and when the average tumor volume reached approximately 50 mm^3^ on day 9, they were intramuscularly injected with mCAP-Fluc, UPA-E6E7, 2UPA-E6E7, or mCAP-E6E7 vaccines (3 μg per mouse, n = 6 for each group), followed by a second dose (3 μg per mouse) one week later (day 16). (B)Tumor volumes from day 0 to day 23 after tumor inoculation. Statistical significance was calculated based on tumor volumes data on day 23. (C) Survival rate of mice from day 0 to day 40 after tumor inoculation. (D) Proportion of E6E7-specific CD8+ T cells in PBMCs one week after the second vaccine dose (IFN-γ-positive T cells after stimulation with E6E7 antigen peptides). (E) Representative flow cytometry plots for (D). (F) Proportion of E6E7-specific CD8+ T cells in the spleen on day 40 (IFN-γ-positive T cells after transfection with E6E7 mRNA). For (B), data are mean (SEM). All other data are mean (SD) for 6 biological replicates. For (C) Kaplan-Meier simple survival analysis was used to calculate the survival rate, and Log-rank (Mantel-Cox) test was used to calculate the statistical significance. For other plots, one-way ANOVA was used to calculate the statistical significance. *P < 0.05 was considered statistically significant. **P < 0.01, ***P < 0.001 and ****P < 0.0001 were considered highly significant. ns, not significant.

PBMCs were harvested one week after the booster dose and restimulated with E6/E7 peptides. FACS results showed high frequencies of IFN-γ⁺ T cells in all E6E7 vaccine groups (mean IFN-γ⁺ CD8⁺ T cells proportions: UPA-E6E7 9.6%; 2UPA-E6E7 15.5%; mCAP-E6E7 13.6%; mCAP-Fluc 0.1%) (Figure 6D, 6E), with 2UPA-E6E7 eliciting the strongest response. On day 40, splenocytes restimulated ex vivo with mCAP-E6E7 mRNA again showed the highest frequency of IFN-γ⁺ CD8⁺ T cells in the 2UPA-E6E7 group, significantly exceeding that of the mCAP-E6E7 cohort (Figure 6F, S7A). Collectively, these results demonstrate that 2UPA-E6E7 LciRNA elicit robust and durable antigen-specific CD8⁺ T-cell responses that correlate with their superior antitumor efficacy.

## DISCUSSION

We engineered a streamlined, cap-independent LciRNA vaccine platform by combining the USUV sfRNA-derived UX1 sequence with a PABP motif to create the UPA protective element, and pairing it with the cap-independent EV-A-S1 IRES. Unlike conventional capped mRNA, incorporation of modified ribonucleotides into LciRNA markedly reduced its expression. In contrast, tandem duplication of UPA (2UPA) yielded an unmodified LciRNA whose *in vivo* stability in mice surpassed that of m1Ψ-modified cap1 mRNA. Mechanistic analysis revealed that UPA not only blocks XRN-1 mediated decay but also recruits the stabilizing RBPs PABPC1, IGF2BP1, and YBX1 to further enhance LciRNA stability. Transcriptome profiling of BMDCs transfected with LciRNA demonstrated potent activation of PRRs signaling, enhanced dendritic cell maturation, and upregulation of antigen-presentation and proinflammatory cytokine expression, indicating a strong intrinsic self-adjuvant effect. Safety evaluation by monitoring body weight, serum ALT and AST levels, and histopathological analysis confirmed that LciRNA vaccine is well tolerated in mice. Finally, in both B16F10 melanoma and HPV-associated tumor models, LciRNA-based vaccines elicited superior immunogenicity and tumor control compared with traditional capped, modified mRNA vaccines.

Comparing with existing mRNA vaccine platforms, LciRNA relies solely on functional RNA sequences—no exogenous proteins, complex processing, or modified ribonucleotides—simplifying production while outperforming conventional linear mRNA in both expression and immunogenicity, thus defining a novel type of mRNA vaccine platform. Previously, several studies also explored uncapped linear RNA designs: Two groups replaced the 5’ cap with a single IRES, yielding an uncapped RNA successfully expressed in cells and *in vivo*^22,23^. However, our results show that uncapped RNA without any protection decays rapidly (Figure 2D, 2E). Similarly, another group proposed a hairpin-protected uncapped RNA encoding the influenza HA antigen, which successfully elicited antibody response^24^. We found that the artificial hairpin sequence did confer modest protection to the uncapped RNA, but its stability remained markedly lower than that of capped mRNA or LciRNA (data not shown). Given that XRN-1 is capable of resolving complex secondary structures^60,61^, simple hairpin structures are unlikely to offer sufficient protection against its degradation. By contrast, viral xrRNAs adopt a precise three-dimensional embrace^31,38^, that blocks the XRN-1 active site, effectively preventing RNA unfolding and degradation. Our findings provide compelling evidence that tandem duplication of UX1 with a PABP motif (2UPA) substantially stabilizes LciRNA *in vivo*, surpassing the performance of capped mRNA. Moving forward, the LciRNA platform can be further optimized by incorporating additional 3’-end protective elements or refining the UPA and IRES sequences to boost functionality while reducing overall transcript length.

Furthermore, LciRNA exhibits intrinsic immunostimulatory activity, making it an ideal tumor vaccine platform. In dendritic cells, LciRNA engages multiple pattern recognition receptors, leading to enhanced antigen presentation and increased chemokine and cytokine secretion, which in turn drive tumor-specific T cell activation and expansion (Figure 7). In a murine melanoma model, LciRNA also augmented intratumoral T cell infiltration, potentially remodeling the tumor immune microenvironment toward an active state. Solid tumors frequently establish immunosuppressive microenvironments characterized by poor immune cell infiltration, which correlates with adverse clinical outcomes^62^. Although combining mRNA vaccines with immune checkpoint inhibitors has yielded promising results in recent trials^4–6^, these strategies depend on capped mRNA with inherently weak innate immunogenicity, restricting their ability to overcome tumor-induced immunosuppression. In contrast, our LciRNA platform leverages intrinsic self-adjuvant activity to provoke strong proinflammatory signaling, remodel the tumor microenvironment, and enhance therapeutic efficacy both as monotherapy and in combination regimens.

**Figure 7.**
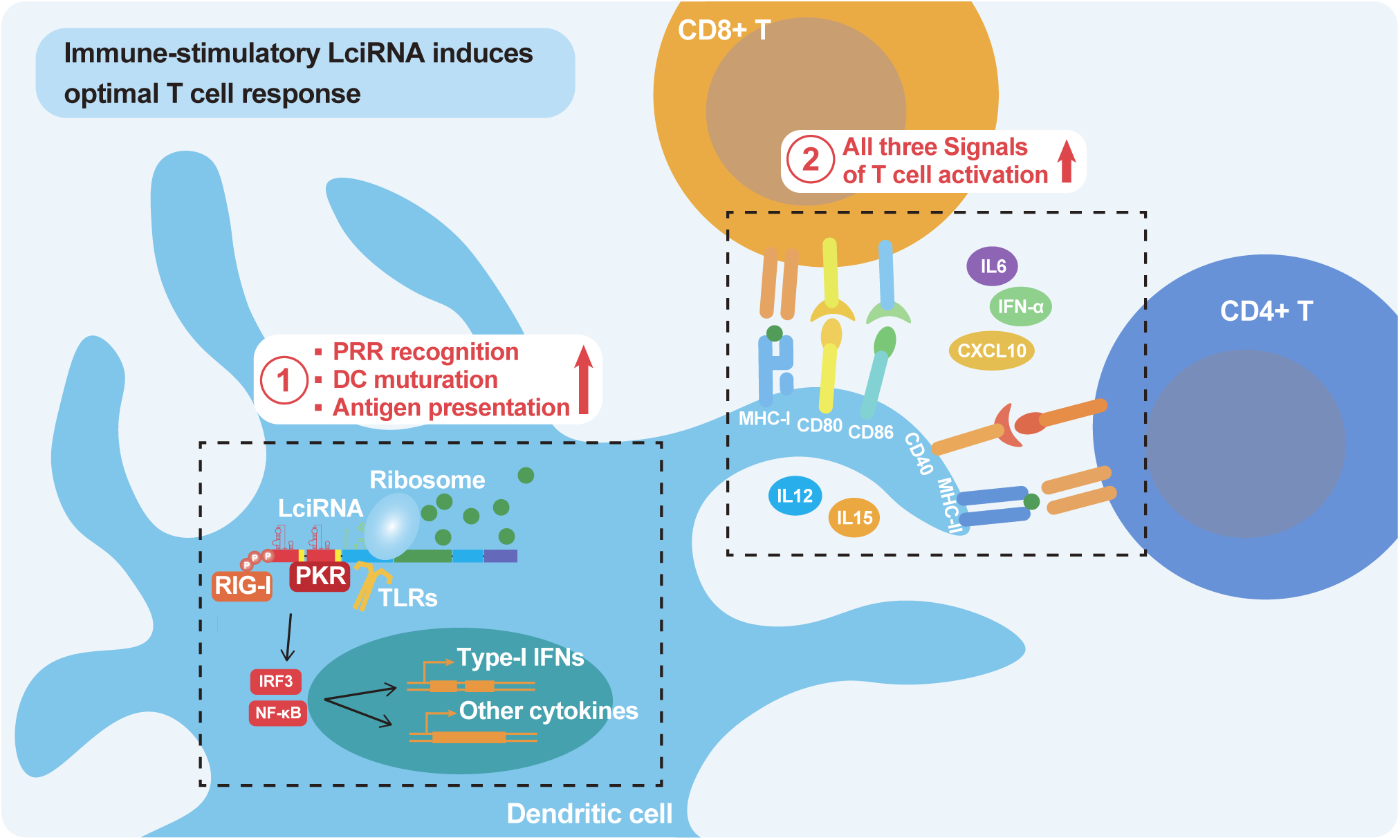
Immunostimulatory LciRNA induces optimal T cell priming. Schematic overview of T cell priming by dendritic cell (DC) after LciRNA cancer vaccine uptake. Upon absorption, LciRNA is recognized by PRRs, such as RIG-I, TLRs and PKR, which activate the expression of Type-I interferon and other inflammatory cytokines and chemokines, such as IL6, IL12, IL15, CXCL10. Then inflammatory signaling induces maturation of DC cells with increased antigen presentation and co-stimulatory protein CD40, CD80, CD86 expression. Meanwhile, relative higher in cell stability of LciRNA insured longer expression of antigenic peptides. As a result, all three signals of T cell activation (antigen presentation, co-stimulatory protein and cytokines) coincide during T cell priming by LciRNA absorbed DC cells. Thus, immunostimulatory LciRNA ensures optimal T cell priming.

The immunostimulatory capacity likely derives from three features of LciRNA: its 5’-triphosphate moiety, which is recognized by RIG-I^52^; the UPA sequence, which binds ILF3 in the cytoplasm and relieves its nuclear suppression of dendritic cell maturation and proinflammatory gene expression in DCs^56^; and the viral-derived IRES, whose extended stem–loop structures could be sensed by PKR and TLRs^53–55^. Together, these features enable multi-PRR engagement and relief of ILF3-mediated immune suppression, robustly inducing downstream immune gene activation (Figure 7). Future mechanistic and translational exploration of LciRNA’s adjuvanticity could maximize its therapeutic potential in cancer immunotherapy.

Nonetheless, this study has several limitations. First, a comprehensive safety assessment of LciRNA will require long-term and higher-dose studies, additional animal species, and more physiological readouts. Second, while we demonstrated specific interactions between the UPA motif and ILF2, ILF3, and PKR, the downstream biological effects are uncharacterized. Notably, ILF3 has established roles in mRNA stability and immune suppression^42,56^, while ILF2 mainly contributes to cell cycle control and apoptosis^44,63,64^. Their association with UPA may influence LciRNA’s immunogenicity and tolerability. Additionally, PKR knockdown enhanced LciRNA translation, suggesting that PKR both restricts expression and promotes immune activation. Future studies must balance these functions to optimize LciRNA’s therapeutic profile.

In summary, LciRNA obviates capping and ribonucleotide modifications, simplifying manufacture and reducing cost while offering enhanced stability and intrinsic adjuvant activity. By engaging diverse PRRs and promoting DC maturation, LciRNA enhances T cell infiltration in “cold” tumors, expanding the therapeutic reach of RNA vaccines. As a fully sequence-encoded, fourth-generation mRNA vaccine platform, LciRNA establishes a new design paradigm for future RNA vaccines.

## MATERIALS AND METHODS

### Molecular cloning

All the LciRNA template sequences were cloned into pUC57 plasmid through seamless cloning. The Cloning Kit for mRNA Template (Takara Bio) was used to construct all the capped mRNA templates. DNA fragments were synthesized by Genewitz (Suzhou, China) and amplified by PCR. ClonExpress II One Step Cloning Kit form Vazyme (Nanjing, China) were used for seamless cloning. All the related sequences are provided in Table S1 and Table S2.

### LciRNA and capped mRNA synthesis

For both types of RNAs, the DNA templates for IVT were first amplified by PCR using 2×AdvanceFast PCR Master Mix (Yeasen, Shanghai) and primers with T7 promoter and 120nt poly(T) (to add 120nt poly(A) tail during transcription). The LciRNAs were synthesized using T7 High Yield RNA Synthesis Kit (Yeasen, Shanghai) without addition of any cap analog or modified ribonucleotides. The modified LciRNAs were synthesized using T7 High Yield RNA Synthesis Kit (Yeasen, Shanghai) with indicated proportion of modified ribonucleotide added (m6A, m1Ψ, or N4-Ac-C from Syngenebio), but without addition of any cap analog. The capped mRNAs were synthesized using T7 High Yield RNA Synthesis Kit (Yeasen, Shanghai) with CleanCap Reagent AG (TriLink) and m1Ψ (Syngenebio) added co-transcriptionally. Both types of RNAs were subsequently treated with DNase I at 37℃ for 20 mins, and purified using GeneJET RNA Purification Kit (Thermofisher). Final RNA products were analyzed by 2% agarose gel or capillary electrophoresis.

### Luciferase reporter assay

For gaussia luciferase assay, cells were cultured in 96 well plate at the density of 5,000 cells per well. The next day, mRNAs were transfected with CALNP mRNA *in vitro* reagent (D-nano, Beijing) at 50ng per well according to manufacturer’s instruction. 24 hours later, 30μL cell culture medium was taken from each well, and luminescence counts were measured using Gaussia Luciferase Reporter Gene Assay Kit (Beyotime) according to manufacturer’s instruction.

### SHAPE-MaP analysis of UX1 structure

The UX1 RNA structure was analyzed by SHAPE-MaP method reported by Smola et al [35] with some modifications as described by our previous study^9^.

### siRNA knockdown

HEK293T cells were cultured in 24 well plates at 100,000 cells per well. siRNAs targeting corresponding proteins were transfected at 30pmol per well with Hieff Trans siRNA/miRNA reagent (Yeasen). 48 hours later, total RNAs of three of the replicate wells were extracted and assayed by RT-qPCR to analysis knockdown efficiency. mRNAs were transfected into the other three replicates, at 200ng per well with TransIT-mRNA Transfection Kit (Mirusbio). 24 hours later, 30μL cell culture medium was taken from each well to measure gaussia luciferase signal. siRNA sequences were synthesized by Hipobio (Huzhou, Zhejiang) and provided in Table S3.

### LNP encapsulation of RNA

All RNAs used in injection of mice were encapsulated by the SM102 LNP mix using GenNano-E0021 cancer vaccine kit (Micro&Nano, Shanghai) according to manufacturer’s instruction. Qubit 2.0 (Invitrogen) and Qubit RNA HS (Invitrogen) were used to determine the RNA concentration in the RNA-LNP vaccine (2% Triton X-100 was added to disrupt the RNA-LNP sample).

### *In vivo* bioluminescence imaging in mice

The Fluc RNA-LNP (5 μg RNA per mouse) was intramuscularly injected into the right thigh of the mice. At indicated time points, 200 μL of D-luciferin potassium salt (Meilun Biology, Dalian) was administered per mouse. Fluorescence imaging was performed and analyzed using the IVIS *in vivo* imaging system (PerkinElmer) within 10–30 min after substrate injection.

### XRN-1 Digestion Assay

For the XRN-1 digestion assay, 2 μg of RNA was first denatured at 70 °C for 2 min and immediately placed on ice. The RNA was then refolded in a 20 μL reaction containing 1× NEBuffer 3 (NEB) and 1 U/μL RNase Inhibitor murine (Vazyme) at 37 °C for 10 min. Subsequently, 1 U of XRN-1 (NEB) and 10 U of RppH (NEB) were added, followed by incubation at 37 °C for 2 h. After digestion, 5 μL of the reaction mixture was mixed with 5 μL of 2× RNA loading buffer, heat-denatured at 95 °C for 1 min, and analyzed alongside an equal amount of undigested RNA using 2% agarose gel electrophoresis.

### RNA pull-down assay

Streptavidin-coated magnetic beads (QuarAcces, Diying, Shanghai) were equilibrated at room temperature for 30 min, washed twice with Solution A (0.1 M NaOH, 0.05 M NaCl in DEPC-treated H_2_O) and once with Solution B (0.05 M NaCl in DEPC-treated H_2_O), then resuspended in Solution B. In parallel, 30 µg of streptavidin-aptamer–tagged RNA was denatured at 56 °C for 5 min in RNase-free water, refolded in annealing buffer (100 mM Tris-HCl pH 7.5, 300 mM NaCl, 3 mM MgCl_2_, 2 U/µL murine RNase inhibitor) at 37 °C for 10 min followed by 5 min at room temperature, and incubated with the pretreated beads for 10 min at room temperature and then rotated for 2 h at 4 °C. HeLa cell lysates were cleared by centrifugation (15 000 × g, 15 min, 4 °C), pre-adsorbed with empty beads for 30 min at 4 °C, and subsequently incubated with the RNA-bound beads in the presence of 1.5 mM MgCl_2_ for 4 h at 4 °C. Beads were washed six times with NT2 buffer (50 mM Tris-HCl pH 7.5, 150 mM NaCl, 1.5 mM MgCl_2_), and proteins were eluted in 1× SDS loading buffer by boiling at 100 °C for 10 min. Finally, eluates were resolved on a 10% SDS-PAGE gel (18 × 16 cm) at 40 V, silver-stained (Beyotime Biotechnology Fast Silver Stain Kit), and differentially visible bands excised for mass spectrometry analysis.

### Western blotting

Pull-down protein samples and 5% Hela cell lysate input were separated on a 10% sodium dodecyl sulfate-polyacrylamide gel and transferred on to a nitrocellulose membrane (Bio-Rad, Hercules, CA, USA). The membrane was next blocked with 5% non-fat milk and incubated with antibodies of indicated proteins or β-Actin antibody. The protein bands were detected using a Tanon 5200 Chemiluminescent Imaging System (Tanon, Shanghai, China) with Omni-ECL reagents (Epizyme Biotech, China). Antibodys used in western blotting were YBX1 (84993-4-RR, Proteintech), PABPC1 (66809-1-Ig, Proteintech), ILF2 (67685-1-Ig, Proteintech), ILF3 (68213-1-Ig, Proteintech), IGF2BP1 (22803-1-AP, Proteintech), EIF2AK2 (66646-1-Ig, Proteintech), DDX3X (67915-1-Ig, Proteintech), HNRNPR (29980-1-AP, Proteintech), β-Actin (66009-1-Ig, Proteintech).

### RNA immunoprecipitation assay

2×5μg UPA-SA aptamer RNA was transfected into two 10cm plates of Hela cells 24 hours before harvesting the cells. Cells were harvested by scraping in 1 mL PBS, and pelleted by centrifugation at 800 ×g for 5 min at 4°C. The cell pellet was lysed in 800 μL RIP lysis buffer supplemented with RNase inhibitor (Vazyme) and protease inhibitor (MedChemExpress), and incubated on ice for 30 min before storage at −80°C. For immunoprecipitation, 50 μL protein G magnetic beads per sample were washed twice with NT2 buffer, incubated with 5 μg target antibody or control IgG in 200 μL NT2 buffer for 1 h at room temperature with rotation, washed again, and resuspended in 500 μL ice-cold NT2 buffer. Cell lysates were cleared by centrifugation (15,000 ×g, 20 min, 4°C), and 100 μL supernatant was incubated overnight at 4°C with antibody-bound beads in 900 μL precipitation buffer (NT2 buffer with 35 μL 0.5 M EDTA and 5 μL RNase inhibitor). Beads were washed six times with NT2 buffer, and 50 μL was reserved for Western blot validation. RNA was extracted using TRIzol (Invitrogen), reverse transcribed, and analyzed by qPCR to assess enrichment. Antibodys used in RIP assay were Rabbit IgG (30000-0-AP, Proteintech), Mouse IgG (B900620, Proteintech), YBX1 (84993-4-RR, Proteintech), PABPC1 (66809-1-Ig, Proteintech), ILF2 (67685-1-Ig, Proteintech), ILF3 (68213-1-Ig, Proteintech), IGF2BP1 (22803-1-AP, Proteintech), EIF2AK2 (66646-1-Ig, Proteintech), DDX3X (67915-1-Ig, Proteintech), HNRNPR (29980-1-AP, Proteintech).

### RNA extraction and RT-qPCR

Total RNA was extracted using TRIzol (Invitrogen) following the manufacturer’s instructions. cDNA was synthesized from using Evo M-MLV Reverse Transcription Kit (Accurate Biotechnology, Huan) following the manufacturer’s instructions. mRNA levels were quantified using SYBR Green Premix Pro Taq HS qPCR Kit (Accurate Biotechnology, Huan). Primer sequences are provided in Table S4. All qPCR reactions were performed on QuantStudio 5 Real-Time PCR System (Applied Biosystems).

### siRNA knockdown

Hela cells were cultured in 24 well plates at 100,000 cells per well. siRNAs targeting corresponding proteins were transfected at 30pmol per well with Hieff Trans siRNA/miRNA reagent (Yeasen). 48 hours later, total RNAs of three of the replicate wells were extracted and assayed by RT-qPCR to analysis knockdown efficiency. mRNAs were transfected into the other three replicates, at 200ng per well using CALNP mRNA *in vitro* reagent (D-nano, Beijing). 24 hours later, 30μL cell culture medium was taken from each well to measure gaussia luciferase signal. siRNA sequences were synthesized by Hipobio (Huzhou, Zhejiang) and provided in Table S4.

### BMDC stimulation asssay

BMDCs were generated with the method reported by Lutz et al. ^66^ For BMDC stimulation assay, BMDCs were cultured in 24 well plates at 250,000 cells per well with mGM-CSF (Peprotech) added to 4ng/mL. 6 hours later, 800ng indicated RNAs or transfection control were transfected into each well using CALNP mRNA *in vitro* reagent (D-nano, Beijing). 24 hours later, total RNAs were extracted and analyzed by RT-qPCR as well as RNA-seq.

### RNA-Seq Analysis

Total RNA from BMDC samples were isolated using TRIzol (Invitrogen) following the manufacturer’s instructions. To eliminate DNA contamination, total RNAs were treated with DNase I (Yeason). The RNA-seq libraries were prepared by using SMARTer Stranded Total RNA-Seq Kit - Pico Input Mammalian (Yeason). Quality control was performed using Qubit (Thermo Fisher Scientific, USA) and Qsep100 (BiOptic, China) before the libraries were sequenced on the BGI DNBSEQ-T7 Sequencing Platform using a 150 bp paired-end run. The RNA-seq data sequencing reads were aligned to the reference genome (Genome Reference Consortium GRCh38) using the spliced read aligner STAR, which was provided with the Ensembl human genome assembly. The gene expression matrix was obtained by featureCounts and be normalized by TPM for downstream analysis.

For data analysis, genes with |log_2_FC| > 1 and adjusted p-value (FDR) < 0.05 were classified as differentially expressed (DEGs). Gene Set Enrichment Analysis (GSEA) was performed using the clusterProfiler v4.2.2 R package. DEGs were ranked by log_2_FC values and tested against the Molecular Signatures Database (MSigDB). Pattern recognition receptor (PRR)-related genes were curated from literature-defined sets encompassing Toll-like receptors (TLRs), NOD-like receptors (NLRs), RIG-I-like receptors (RLRs), and C-type lectins (CLRs). Heatmaps were generated using the pheatmap v1.0.12 R package: DEG expression values were transformed to z-scores by row, and visualized with diverging color gradients.

### Animal model

Male C57BL/6 and BALB/c mice aged between 6-8 weeks were purchased from Biocytogen (Beijing, China).

For in vivo bioluminescence assay, BALB/c mice were injected with 100μL Fluc RNA-LNPs (5μg RNA per mouse) via intramuscular injection to the right thigh. At the indicated time points, 200 μL D–luciferin potassium salt solution (15mg/mL) was injected to each mouse through intraperitoneal injection. In vivo luminescence signal was measured using an IVIS Spectrum system (PerkinElmer) 10 min later. Data were analyzed using Living image software.

For serum ALT and AST evaluation, mouse blood was collected at the indicated time points, and kept at room temperature for 60 min for coagulation. Serum samples were collected through centrifugation at 3000 rpm/min under 4℃. Then, serum samples were sent to Fudan University Laboratory Animal Center for analysis using ADVIA XPT (SIEMENS). For HE-staining of mouse tissue, after euthanizing the mice, organ samples were collected, fixed and paraffin-embedded, sectioned, and stained for H&E (Servicebio).

For the murine melanoma model, 1×10^5^ B16F10-OVA cells in 200ul RPMI 1640 medium (without fetal bovine serum) were subcutaneously injected into the right flank of C57BL/6N mice. For the HPV associated TC-1 model, 1×10^5^ TC-1 cells in 200ul RPMI 1640 medium (without fetal bovine serum) were subcutaneously injected into the right flank of C57BL/6N mice. Mice body weight and tumors were measured twice every week. Tumor volumes were determined by using the formula volume = (length × width^2^)/2 (L: largest diameter of the tumor; W: smallest diameter of the tumor). For immunization, the mice were vaccinated with 100μL RNA-LNPs (3μg RNA per dose) via intramuscular injection to the right thigh. All experiments were performed in accordance with protocols approved by Fudan University Experimental Animal Care Commission.

### Flow cytometry analysis

For OVA (SIINFEKL) Tetramer staining, PBMCs was separated by Mouse 1× Lymphocyte Separation Medium (DAKEWE, Shenzhen) according to the manufacturer’s instruction. Cells were then stained with Fixable Viability Dye eFluor 450 (65-0863-14, eBioscience), anti-CD3 (APC-65077, Proteintech), anti-CD8a (FITC-65069, Proteintech), H-2Kb&B2M&OVA (SIINFEKL) Tetramer (PE, ACROBiosystems), and analyzed by CytoFLEX Flow Cytometer (Beckman).

For intratumoral T cell staining, tumor tissues were dissected into 1–2 mm³ fragments in ice-cold RPMI 1640 medium (10% FBS). Samples were digested in a cocktail of 1 mg/mL collagenase IV (Yeasen) and 100 U/mL DNase I (Yeasen) for 30 min at 37°C with constant agitation (200 rpm). Cells were then stained with Fixable Viability Dye eFluor 450 (65-0863-14, eBioscience), anti-CD3 (APC-65077, Proteintech), anti-CD8a (FITC-65069, Proteintech), anti-mouse CD4 (PE-CY7, Biolegend), H-2Kb&B2M&OVA (SIINFEKL) Tetramer (PE, ACROBiosystems), and analyzed by CytoFLEX Flow Cytometer (Beckman).

For intra-cellular IFNγ staining, PBMCs or splenocytes was separated by Mouse 1× Lymphocyte Separation Medium (DAKEWE, Shenzhen) according to the manufacturer’s instruction. Cells were then stained with Fixable Viability Dye eFluor 450 (65-0863-14, eBioscience), anti-CD3 (APC-65077, Proteintech), anti-CD8a at 4℃ for 30min. Subsequently, cells were fixed and permeabilized with Intracellular Fixation and Permeabilization Buffer Set (eBioscience) according to the manufacturer’s instruction, and stained with anti-IFNγ (PE, Biolegend) at room temperature for 20min, and analyzed by CytoFLEX Flow Cytometer (Beckman).

### Statistical analysis

All data in this study were analyzed via Graphpad Prism and presented as mean (SD) unless otherwise indicated. Two-tailed unpaired Student’s t-test was used to compare two groups. One-way ANOVA tests were used to compare more than two groups. For survival curves, the data was performed via Kaplan-Meier analysis. *P < 0.05 was considered statistically significant. **P < 0.01, ***P < 0.001 and ****P < 0.0001 were considered highly significant. ns, not significant.

## DATA AND CODE AVAILABILITY

The authors declare that all data supporting the findings of this study are available within the article and its supplemental files or are available from the authors upon request.

## ACKNOWLEDGMENTS

This work was supported by grants from National Key Research and Development Project of China (2021YFA1300500), National Natural Science Foundation of China (82272625), and Fujian provincial health technology project (2024ZD01004).

## AUTHOR CONTRIBUTIONS

S.H., H.Y., Z.H. and Z.F. designed the study; H.Y. performed the experiments and analyzed the data; Y.Y., C.L., and Y.W. helped with some of the experiments; P.L. helped with the RNA-seq data analysis; H.Y., S.H. and Z.H. wrote and revised the manuscript. All authors have read and agreed to the published version of the manuscript.

## ABBREVIATIONS

mRNA: messenger RNA
LciRNA: Linear cap-independent RNA
circRNA: circular RNA
saRNA: self-amplifying mRNA
dsRNA: double-stranded RNA
IRES: internal ribosome entry site
DC: dendritic cells
RBPs: RNA-binding proteins
sfRNA: subgenomic flaviviral RNA
PABP: poly(A)-binding protein
GLuc: Gaussia luciferase
FLuc: firefly luciferase
EV-A: *Enterovirus* A
xrRNA: exoribonuclease-resistant RNA
m1Ψ: 1-methyl-pseudouridine
m6A: N6-methyladenosine
N4-Ac-C: N4-acetylcytidine
UTR: untranslated sequences
poly(A): polyadenylate
SHAPE-MaP: selective 2’hydroxyl acylation analyzed by primer extension and mutational profiling
UPA: UX1 plus PABP motif
RIP: RNA immunoprecipitation
BMDC: bone marrow-derived dendritic cell
TLRs: Toll-like receptors
PRRs: pattern recognition receptors
GSEA: gene set enrichment analysis
ALT: alanine aminotransferase
AST: aspartate aminotransferase
H&E: hematoxylin and eosin
PBMCs: peripheral blood mononuclear cells
OVA: ovalbumin
HPV: human papillomavirus
RFS: recurrence-free survival

---

**Figure S1. RNAfold predicted xrRNA and dumbbell structures of CFAV, DENV, USUV, YFV and ZIKV.**

Minimum free energy (MFE) structures were predicted with RNA energy parameters of Andronescu model chosen. The structures were visualized by Forna.

**Figure S2. UX1 SHAPE-MaP reactivity profile and top ranked proteins in mass spectrometry result.**

(A) Bar plot of SHAPE-MaP reactivity profile of UX1. Reactivities below 0.4 are colored black, reactivities between 0.4 and 0.85 are shown in orange and reactivities above 0.85 are shown in red. (B) Top ranked proteins in mass spectrometry result. The red box indicates proteins distributed in the cytoplasm.

**Figure S3. Heatmaps of normalized mRNA expression levels of genes upregulated in PRR induced DC maturation, and genes of T cell co-stimulatory protein, inflammatory cytokines and chemokines.** (A) Heatmap of normalized mRNA expression levels of genes upregulated in PRR induced DC maturation. (B) Heatmap of normalized mRNA expression levels of genes of T cell co-stimulatory protein, inflammatory cytokines and chemokines. Rows represent individual genes and columns represent experimental samples (n=3 biological replicates per group). TPM values were scaled by row to standardize gene-wise expression patterns, with color intensity indicating relative expression levels (blue, low expression; red, high expression).

**Figure S4. Body weight means, tumor volumes of individual mouse and representative flow cytometry plots of the B16F10-OVA treatment model in Figure 5.**

(A) Mean body weight of each group from day 0 to day 20. Data are mean (SD) of n=6 biological replicates per group. (B) Tumor volumes of individual mouse from day 0 to day 20. (C) Representative flow cytometry plots of Figure 5F.

**Figure S5. Gating information of Figure 5.**

(A), (B) and (C) are gating information of PBMCs (Figure 5D), spleens (Figure 5F), tumors (Figure 5G, 5H and 5I).

**Figure S6. Body weight means, tumor volumes of individual mouse and representative flow cytometry plots of the B16F10-OVA treatment model in Figure 5.**

(A) Mean body weight of each group from day 0 to day 23. Data are mean (SD) of n=6 biological replicates per group. (B) Photographs of mice tumors on day 23. (C) Tumor volumes of individual mouse from day 0 to day 23.

**Figure S7. Representative flow cytometry plots and gating information of Figure 6.**

(A) Representative flow cytometry plots of Figure 6F. (B) Gating information of PBMCs in Figure 6D. (C) Gating information of PBMCs in Figure 6F.

**Table S1. xrRNA, Dumbbell, IRES, UPA and UTR sequences**

**Table S2. Aptamer, Reporter and Vaccine sequences**

**Table S3. siRNA**

**Table S4. qPCR primers**

